# Greylag geese (Anser anser) initiate group flight based on the number and identities of callers

**DOI:** 10.64898/2026.10.26.761644

**Authors:** Dina A. A. Mostafa, Alan G. McElligott, Nicolas M. Adreani, Elodie F. Briefer

## Abstract

Social animals need to maintain cohesion when they move, and movement decisions are influenced by social organization and ecological or individual needs. To maintain synchrony during collective movement, the use of signals, including vocal signals, is often necessary. Greylag geese are gregarious and vocal before and during flight. We observed instances of naturally occurring flight initiation attempts in a free-living population of greylag geese (n = 133 birds) to investigate the decision-making process in this species, including the role vocalisations play in flight decisions. We recorded the number of participants in flight-initiation attempts, their vocal behaviour (using two call types specific to flight initiation signalling: Locomotion and Pre-flight calls), the outcome of those attempts, in addition to the resulting flying group size and composition. We found that the identities and number of flock members joining in the calling chorus predicted both the probability of successful flight and the resulting flying group size. The number of Locomotion callers was associated with an increased probability of success, as well as larger flying group size, including both family and non-family fliers. The number of Pre-flight callers was also associated with a higher probability of successful flight, but its effect on the increase in flying group size was marginal. Successful departures were more probable when the family members of the initiator responded vocally. Overall, our findings are consistent with a system where the decision to fly is made individually using two distinct vocal signals to estimate the number of ‘voters’ and (at least) the identities of social partners and progeny. In species where temporal and spatial group cohesion varies both seasonally and diurnally, individuals’ priorities appear to be maintaining proximity to mates and family members, followed by maximizing the size of flight groups.

## Introduction

Group living increases the fitness of social animals through shared predator vigilance, by providing parental care and continued mate guarding (Krebs & Dawkins, 1984; Clark & Mangle, 1986; Covas & Griesser, 2007; Black & Owen, 1989), or by gaining information about foraging sites (Valone & Templeton, 2002). As a result, maintaining the cohesion of social groups has a direct positive effect on lifelong individual fitness and reproductive success (Black & Owen, 1989; Mcfarland et al., 2017; Silk et al., 2009; Cameron et al., 2009).

Group cohesion requires synchronicity in the decision to move, which can conflict with the individual needs of some group members; because individuals might vary in their physiological needs of foraging and rest, for example, the motivation to switch from one activity to another that involves movement can differ within pairs, family, and among group members (Conradt & Roper, 2000; Valone & Templeton, 2002). Since group movement increases the risks of separation, it becomes essential for social affiliates and group members to reach a decision about the timing of movement together, and to advertise their motivation to move if group integrity is to be maintained (Sueur et al., 2011; Xie et al., 2024). Therefore, animal groups are under pressure to evolve signalling mechanisms by which all members share information regarding movement readiness, and means by which the mismatch in motivation and readiness can be resolved. These selective pressures have given rise to various types of collective decision-making and associated signalling patterns, and resulted in synchronous movement often requiring non-independent decisions based on public information (Farine, 2022; Petit & Bon, 2010; Strandburg-Peshkin et al., 2015).

The type of collective decisions depends on which individuals contributes to the decision, and this varies between systems according to some of their key characteristics, such as social dominance hierarchy, the spatiotemporal cohesion of different units, as well as mating and pairing systems (Prox & Farine, 2020b). For instance, movement decisions in species with linear hierarchies are not shared among the group, but rather made by one dominant individual (i.e. ‘despotic’; Nellissen et al., 2024). The individual with the highest rank signals the information about eminent movement, and group members follow even when consensus comes with an adaptive cost (Conradt & Roper, 2003; King et al., 2008). Alternatively, collective decisions can be shared (i.e. ‘democratic’) when reached by multiple individuals. This type of decision is typically seen in groups with non-linear, changing hierarchies. The decision in those social systems is either made by individuals belonging to a particular caste in the case of eusocial insects (Seeley & Visscher, 2004), or equally by all group members, as observed in many primate and bird species (Black, 1988; Boinski & Campbell, 1995; Lamprecht, 1992). Participants cast their ‘votes’ via the use of specialised signals, and movement commences once a particular threshold is reached (Bousquet et al., 2011). Despite the overall distinction between these two opposite types of group decision making found in the literature, a clear distinction is not always found in biological systems, which necessitates careful consideration when interpreting how different animal groups reach decisions (Pyritz et al., 2011)

Social organisation, defined as the size and composition of a social unit (Prox & Farine, 2020a), also influences the cohesion of the group under movement; in groups with high spatiotemporal integrity, movement decisions are reached by consensus, which involves one of two mutually exclusive actions. This is the case of colony relocation of eusocial insects (Seeley & Buhrman, 1999), in which the group moves as one unit, including the individuals that did not participate in making the movement decision (Conradt & Roper, 2007). On the other end of the continuum, we find animal groups with fluid social structure, where smaller social units coalesce to form larger groups that then break up over time and space (Farine & Whitehead, 2015; Fryxell & Berdahl, 2018; Kerth et al., 2006; Sigaud et al., 2017). In these systems, the individual makes the decision after integrating the publicly available information relevant to the readiness of other group members, with knowledge of own needs, to make a ‘combined decision’ about moving with others or staying behind (Deneubourg et al., 2002; Valone & Templeton, 2002). Combined decisions are not independent on the group level, since they rely on information shared by many members (Sperber et al., 2019). Pairing systems and social bonds play an important role in social cohesion and subsequently in movement decisions, as these determine the primary social unit in group living animals, and maintaining the cohesion of these units maximises the adaptive advantage of social living.

Which pieces of information are integrated into making a collective decision depends therefore on the degree of ‘democracy’ involved and the social organisation of the system (List, 2004). Signal integration plays an essential part in reaching collective decisions, regardless of how that is structured. When considering individual votes, we find that in many species, even when the decisions are shared, not all ‘votes’ are of the same weight, and the participation of particular individuals results in higher rates of movement initiation success (Walker et al., 2017; Ward et al., 2008). Likewise, for combined decisions, the votes of social partners have more weight compared to those of other familiar conspecifics (Griffiths & Magurran, 1999; Sueur, Deneubourg, et al., 2011). Other parameters contribute to how signals are integrated to reach a decision, including the temporal aspect of signal accumulation or decay, which in combination with the number and weight of signals determine how a response threshold is reached (Wang et al., 2015). Group state transition from one state to another is therefore a complex process that depends on all of the above. As a result we see decision rules where a consensus reached via a simple quorum where all signals have equal weight (Franks et al., 2003), others where quorum involves identity weighted signals (Marshall et al., 2017), or rules of signal accumulation and temporal integration, where successive signals or their decay accumulate to contribute towards the decision state (Detrain & Deneubourg, 2008; Kietzman & Visscher, 2015; Seeley et al., 2012). These and many other rules contribute towards the collective decision mechanism, and their contribution require examining before a complete model is synthesised for each system (Conradt, 2012).

Agreement to move as a group is therefore reached by sharing information about the timing and direction of movement via the use and perception of public information delivered through signals and/or cues (Larissa Conradt & Roper, 2000; King, 2010; King & Cowlishaw, 2009; List, 2004). Communicating movement decisions happens via a multitude of signalling modalities, including visual (Black, 1988; Raveling, 1969), acoustic (Poole et al., 1988; Stewart & Harcourt, 1994), and other modalities such as body orientation and or combination of two or more modalities (Xie et al., 2026). In environments where obstacles, distance or crowding can impede certain modes of signalling such as vision, acoustic signals can help convey group wide information instantaneously (Mouterde et al., 2014). Therefore, movement coordination via specialised vocal signals has evolved and spread in animals of all taxa (Xie et al., 2024). Despite the number of studies on the topic being skewed in favour of mammals, particularly apes, movement vocal signals have been observed in a wide variety of other species including birds and insects (Xie et al., 2024). In some species, vocal signals are used to both initiate and direct the group movement (Boinski, 1993; Bousquet et al., 2011), while in others, they are used to reach the decision to move only (Seltmann et al., 2013; Sperber et al., 2017). Vocal signals are particularly useful as they can encode crucial information about a the caller’s attributes (Galeotti et al., 1997; Mager et al., 2007), motivation (Briefer et al., 2019; Huber et al., 2017), and identity (Keen et al., 2013; Węgrzyn & Leniowski, 2020), rendering them particularly suitable for delivering complex array of information (Briefer, 2020; Charlton et al., 2020). The information content pertaining to the caller(s) (e.g. kinship or dominance rank) can, in many contexts, influence the response of the receivers according to their relationship and past interactions with the caller, thus allowing for dynamic social communication (Blumstein et al., 2004; Trillmich et al., 2004).

The greylag goose (*Anser anser*) is a social species that exhibits varying degrees of cohesion in movement and association (Hemetsberger et al., 2013; Lorenz, 1988). Adult geese form long-lasting pair bonds, and parents provide parental care that typically lasts from one breeding season to the next, and may continue for an additional year in case the pair does not produce new offspring (Lorenz, 1988). Preserving family (parent and offspring) cohesion is therefore of great importance for sustained parental care during the first year of the goose’s life, and for mate guarding during the breeding season (Frigerio et al., 2001; Weiß et al., 2011). At higher levels of organisation, flocks are at their most cohesive form during the autumn and winter, and by spring, the birds gradually split from the flock to form pairs and start nesting. Pairs and families gradually coalesce following the fledging of young and adult moult towards the following autumn (Hemetsberger et al., 2013; Lorenz, 1988). Before take-off, individual geese perform a ritualised signalling flight initiation behaviour that combines vocal and motor signalling combined with body orientation (Schmitt, 1991). Similar flight initiation behaviour is also described in closely related species, such as the Canada geese (Raveling, 1969) and whooper swans (Black, 1988). Flight seems to occur once a threshold of excitement or number of callers has been reached. Although the exact condition required for flight initiation in greylag geese has not been quantified, other species are known to initiate group movement either by reaching a certain number or percentage of participants (Franks et al., 2003), or when signalling rate increases to a given threshold (Stewart & Harcourt, 1994).

We investigated the flight initiation vocal behaviour in greylag geese to understand how the choice of flight is reached by individuals of a particular sub-group. The main components of flight initiation signalling are conserved in many species of the swan and geese subfamily (Anserinae) including greylag geese (Cramp & Simmons, 1977; Schmitt, 1991), Canada geese *Branta canadensis* (Raveling, 1969; Black & Barrow, 1985), barnacle geese *Branta leucopsis* (Black, Prop, & Larsson, 2014), whooper Cygnus Cygnus and *Bewick’s swans Cygnus* columbianus, as well as mute swans *Cygnus olor*, (Black, 1988). The stereotyped flight initiation signalling of greylag geese is comprised of a vigilant posture, orientation in the direction of flight, lateral headshakes and two types of calls (Schmitt, 1991), which we provisionally labelled ‘Locomotion call’ and ‘Pre-flight call’. The Locomotion call (Cramp & Simmons, 1977) – also earlier called ‘departure call’ (Lorenz, 1988), is a medium-range (Sound Pressure Level mean @ 1 m = 77.1 dB, N = 31), repeated cackling made of multiple similar elements, typically with an unvarying repetition rate. This call type expresses motivation for movement in general, either on foot, swimming, or flying (Lorenz, 1988). The second call, the Pre-flight call (also earlier called ‘go-away call‘ Schmitt, 1991), is a louder, long-range (SPL mean @ 1 m = 81.7 dB, N = 36), monosyllabic call that is emitted repeatedly when the apparent motivation to fly is strong, typically for few seconds before take-off (Figure. 1). The two call types encode both caller identity and sex (Mostafa & Briefer, 2026), potentially allowing receivers to individually identify callers when visual identification is impeded.

**Figure 1.**
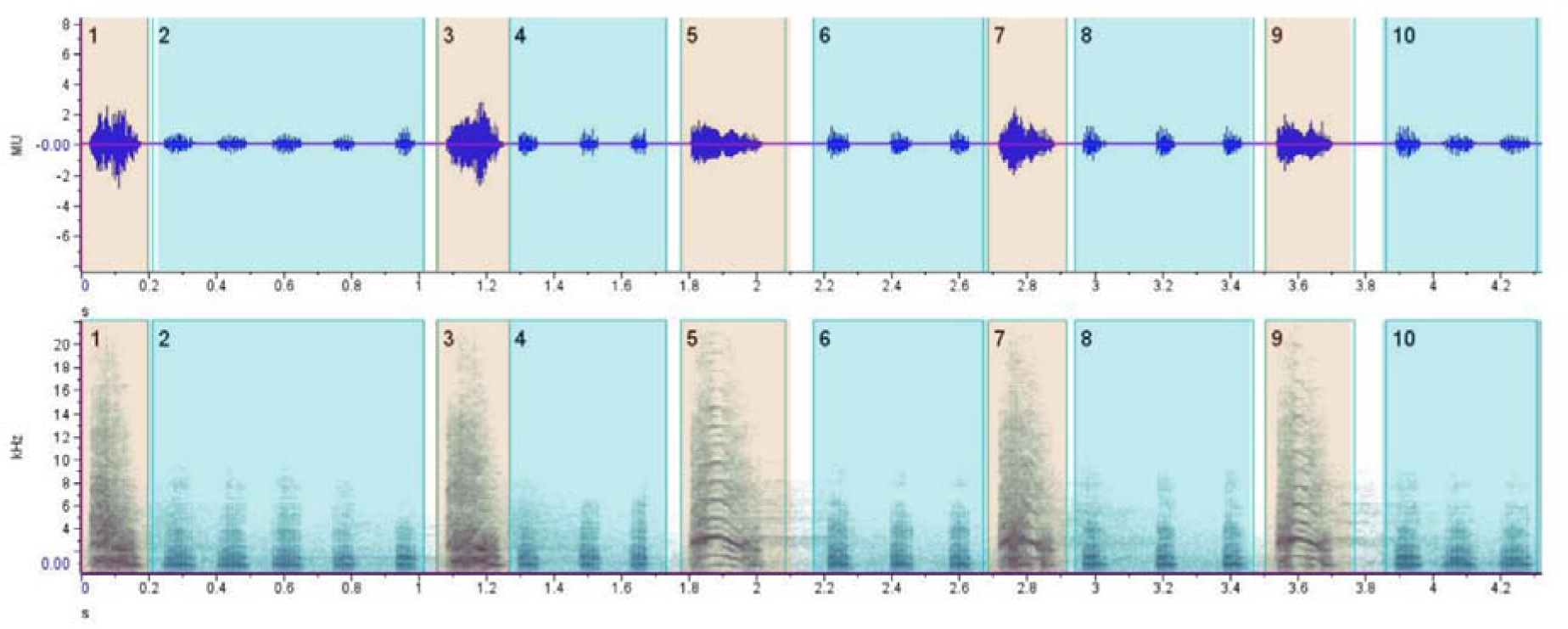
Spectrogram of a Pre-flight calling bout emitted by two individuals (bonded pair). Locomotion calls are emitted here by the female in bouts (even numbers) characterised by rapid repititon, while Pre-flight calls (odd numbers) are emitted here by the male at relatively longer intervals. (Image made with RavenPro) Alt text: Graphical representation of a natural patterns of flight initiation call production in greylag geese, using two different call types. The upper panel represent amplitude change over time, while the lower panel represents frequency change over time.

By observing the vocal behaviour of the participants in the flight initiation signalling, we investigated which variables contribute to the group flight decision of greylag geese. We observed the spontaneous flight initiation attempts of the study population, and recorded the identities of the flight initiating bird as well as those of every bird participating in the flight initiation calling. We also noted the numbers of participants in calling, and whether the attempt resulted in flight. First, we hypothesised that successful group departures always involve acoustic signalling. Vocal signals are often used to coordinate the timing of movement (reviewed in Liao et al., 2024), and we therefore predicted that all flight initiation attempts would be preceded by at least one of the two flight initiation call types described in the literature (i.e. Locomotion and Pre-flight calls: Figure 1). Second, we hypothesised that individual geese prioritise the cohesion of pair and family units during movement, due to the durable social bonds of pair and family described in the species. We therefore predicted that geese would participate in flights only when close social associates express readiness to move vocally. Third, we hypothesised that geese prefer to fly in larger groups. Due to the documented pattern of increased flocking in the species outside of the breeding season, we predicted larger flight groups to also persist during movement, and that the individual pair or family units would seek to join other flock-mates in flight groups, resulting in larger flight group sizes with an increase in number of vocal participants. Finally, because geese do not organise in social groups with linear hierarchies, we hypothesised that their collective movement decision-making is ‘democratic’ and ‘combined’, which involves an equal participation of all group members. As a result, we predicted that the number of callers joining in the flight initiation vocal signalling would positively correlate with the probability of successful flight and increased flying group sizes. Understanding how greylag geese use vocal cues and social information to coordinate departures contributes to our broader understanding of the mechanisms that enable collective movement in animal societies.

## Methods

### Ethical note

The observation of birds was carried out under the license number 66.006/0026-WF/V/3b/2014 by the Austrian Federal Ministry for Science and Research. This study was conducted in compliance with the Austrian Federal Animal Protection Act (TSchG). The banding of the study population is part of a long-term monitoring protocol carried out by the team at the Konrad Lorenz Research Centre (KLF) and the University of Vienna. The study population is habituated to the presence of human observers, and the researcher in this study (DM) conducted daily flock observations for three months before the start of data collection to further habituate the birds. All the data for this study were collected in a non-invasive manner and with no interference with the naturally occurring activities of the birds. The species boasts healthy numbers in Austrian and European wild population (Fox and Leafloor, 2018), and IUCN Red List Category of the species is Least Concern (Birdlife International, 2018).

### Study population

The observations for this study were conducted at the Konrad Lorenz Research Centre (KLF) site in Grünau im Almtal (47_48050.500N 13_56051.000E), a core facility of the University of Vienna, between September 15th and November 9th 2021 for a total of 28 days. We observed a non-migratory, free-living, well-habituated population of Greylag geese established in 1935 by Konrad Lorenz from a stock of eggs that was hand-raised, and all the subsequent cohorts have been under observation since then for the purposes of studying the different aspects of the species behaviour. All birds are individually marked for ease of identification, sexed upon maturation and pair formation through behaviour, and the population pedigree information has been collected since the establishment of the population. The KLF population offers a unique opportunity for studying geese behaviour due multiple characteristics; since its inception from hand-raised individuals, the population has been habituated to continued human presence, and being non-migratory, it spends all year in the valley, making the study of all its life history stages possible. The minimal interference in the upkeep of the birds contributes towards preserving their natural behaviour. The data for this study were gathered during morning and evening group movement events; the birds are provisioned with food of higher nutritional value (grains and grass pellets) compared to the naturally foraged grass, in order to ensure their habitual return to the study site and habituation to human presence. Feeding takes place twice daily, early morning and late afternoon, making their movement to and from the feeding site at KLF predictable and suitable for studying. The flock was also observed at the northern pond of the neighbouring Cumberland Wildpark in the evenings, where they congregate every day before flying back to the feeding site for the evening feeding (Figure 2).

**Figure 2.**
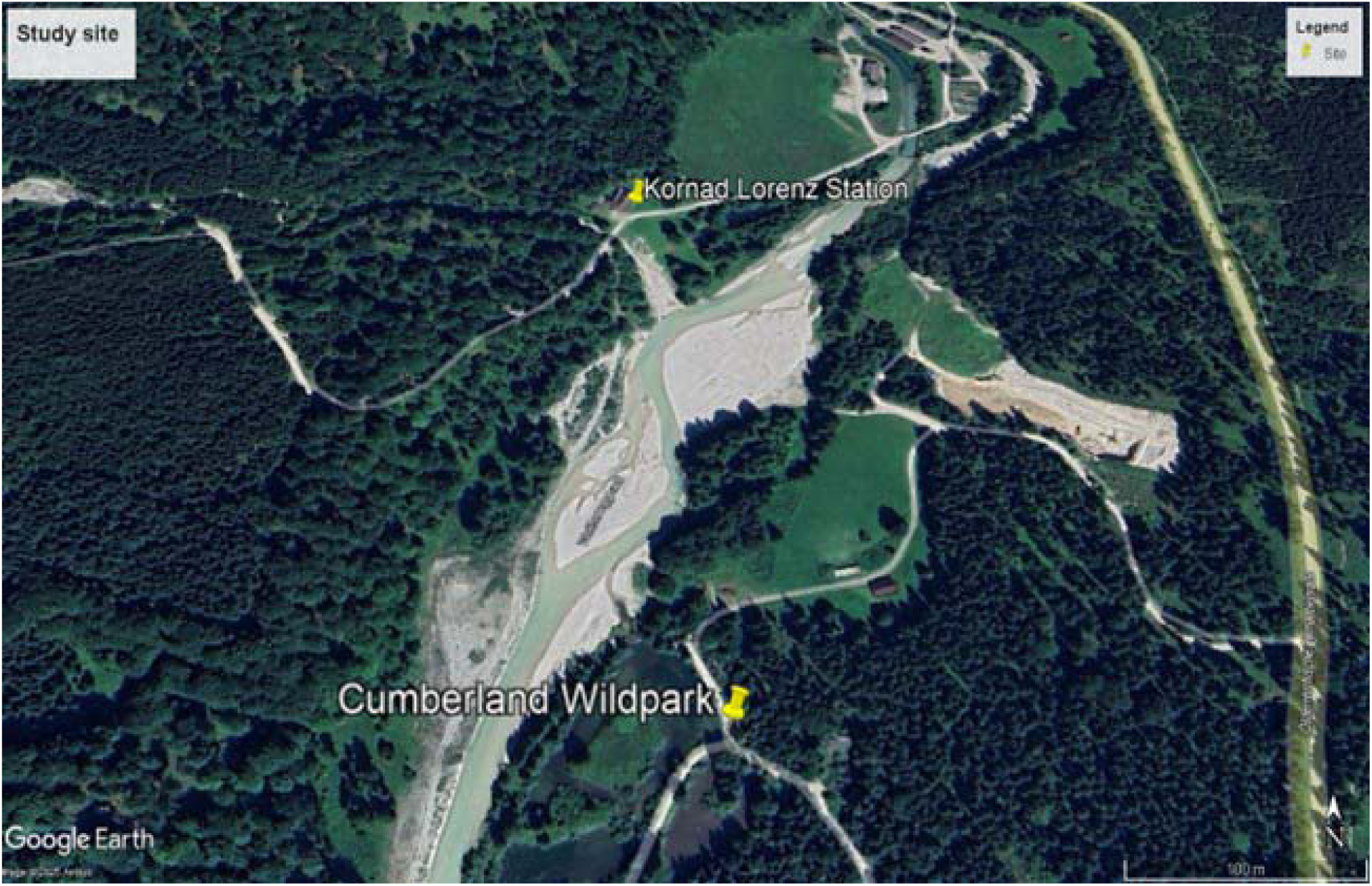
Map of the two observation sites. The Konrad Lorenz Station is the old site of the research centre where the population was established and has since been fed. The site at Cumberland Wildpark is the northern pond where the flock gathers in the evening before flying back to the station for feeding. Image created with Google Earth pro. Alt text: Map representing the study area with the two data collection sites marked showing the distance and trajectory of bird flight between the two.

Observations began right after the young had fully fledged their flight feathers, at around five months of age, and started flying with their parents, which coincides with the completion of the yearly moult by the adults by the end of summer. At that time, the population consisted of 133 individually-marked geese (66 males, 43 females, 24 un-sexed juveniles). The patterns of flock cohesion and movement change seasonally; during the breeding season, breeding pairs break off from the flock, and roost or feed at the nest site. Once the young have hatched, families rejoin the flock but do not fly with it until the young have fledged. The non-breeders form a more cohesive unit that move and feed in smaller units, but roost together at night, and moult in the same location late summer. The autumn – when our data were collected - corresponds to when migratory restlessness would be observed in a wild migratory population, and when increased short distance movement is typically observed in this non-migratory population (Hemetsberger et al., 2013).

### Data collection

Flock movement behaviour commences when the motivation to fly increases, any goose starts performing the ritualised flight initiation behaviour until joined by others, forming multiple flying groups. This process repeats, with different individuals of either sex attempting to initiate flight either successfully or not, until the whole flock had left.

Flight initiation attempts were sampled ad libitum, and a departure attempt observation was considered to have started whenever an individual spontaneously began Locomotion calling and head shaking while moving or facing the direction of flight with vigilant posture. The focal bird was observed while signalling continued and until the trial ended either with a successful departure of one or more individuals (defined as successful flights), or with signalling being aborted for 30 seconds (defined as aborted attempts). This 30 second threshold was chosen because pilot trials showed that all birds participating changed behaviour and returned to feeding or other activities within 30 seconds. The 30 second interval was not implemented for consecutive successful group departures - instead we considered a new attempt to have been started once the previous flying group was entirely out of sight. The following focal bird was chosen and observed from the moment it started the stereotypical flight initiation signalling.

While the focal bird was signalling, the trial was video recorded using a Sony Handycam FDR AX53 with a high-quality mounted microphone (Røde Videomic Pro+, Australia; Freq. range: 40 Hz – 20 kHz, Max SPL: 134 dB). Additional information visible outside the video frame (e.g. identities of other vocalizing individuals) were spoken onto the recording. All spoken observations, including focal and responders’ identities, number of fliers and their identities, duration of signalling, etc., were later confirmed from the videos, which were all captured and saved in 4k resolution, and viewed using commercial media player (VLC media player) with a zoom function and a high-resolution display.

The total number of observed flight initiation attempts was 651. Because flight initiation could be attempted by any individual in the flock, the observer changed her position within the flock following each successful flight to avoid sampling bias. If no attempt was made within the vicinity of the observer, she moved towards the closest signalling bird once a new attempt commenced. Most individuals were within hearing range and easily observed with binoculars (80-90% of birds rested close to each other); the KLF site is an open field with an area of approximately 68 m^2^, and birds were always within 35-50 m of the observer. The pond site at Cumberland Wildpark is an area of approximately 250 m^2^ excluding water. Here, birds that could be used as focals were restricted to those situated on land and within 50 m from the observer. The identities of birds signalling while swimming could not be determined and were later excluded from the analysis, accounting for 50 out of the 651 observations.

We extracted nine variables characterising each departure event (Table 1) designed to address our hypotheses as follows; the number of birds responding with Pre-flight calls (variable 1), and the number of birds responding with Locomotion calls (variable 2), were scored to investigate the idea of a quorum-based collective decision (i.e. the more responders using one or both call types, the greater the probability of success or bigger flying groups). The vocal response of all the family members of the focal (variable 3), was scored to investigate whether geese prioritised the cohesion of their primary social group when making the decision to fly; this variable was scored as ‘Yes’ if all family members (mate and offspring in the case of parents with dependent juveniles, and only the mate in the case of pair bonded birds) responded, and ‘No’ if not all of them responded. For unpaired individuals (26 observations), family consisted of all social associate (i.e. parents and all brood siblings for dependent offspring, and homosocial partners for male-male pairs; 21 observations). By contrast, if the unpaired bird had no social associates (i.e. sub-adults and widowed males; 5 observations), this parameter was always scored as ‘No’. The sex of the focal (variable 4) was scored to investigate whether there were sex-related differences in initiating a successful flight and in recruitment of others for flight. Breeding status (the presence or absence of juveniles that parents provide with parental care; variable 5) was scored to investigate whether parents and non-parents differed in initiating successful flight and recruiting others. The identity of the focal (variable 6) was scored to establish family relationships between initiator and responders, and to control for repeated measures of the same individual. Finally, Session (variable 7; a unique identifier for each observation session) was scored for the investigation of identifiable random intercepts and slopes, and to account for repeated measures.

**Table 1.** Behavioural and demographic variables extracted from the video observations of the spontaneous flight initiation attempts of greylag geese. The variables listed represent the hypothesised causal elements influencing the flight initiation process (1 – 5), sampling repeated measures (6 &7), and outcomes (8 & 9).

| No. | Variable | Description |
| --- | --- | --- |
| 1 | Pre-flight call responders' number | The number and identity of flock-mates responding by emitting Pre-flight calls during the entire trial (Count). |
| 2 | Locomotion call responders' number | The number and identity of flock-mates responding by emitting Locomotion calls during the entire trial (Count). |
| 3 | All family vocal response (yes / no) | Whether the mate (in the case of pairs) or all family members (mate and all progeny in the case of families, or parents, all brood siblings, and homosocial partners for unpaired individuals) of the focal bird participated in vocal signalling during the trial (Binary). |
| 4 | Sex | The sex of the focal bird (Binary). |
| 5 | Breeding status | Whether the focal bird had juveniles they provided with parental care at the time of data collection (Binary). |
| 6 | Focal | The identity of the first bird to signal (ID). |
| 7 | Session | A unique value for each record of flock movement on the wing that spanned all the groups until no bird was left on site on the particular day and time (date + morning/evening), (Integer). |
| 8 | Non family recruited | The number of individuals joining in the group that are neither mates of the focal, nor offspring receiving parental care (Count). |
| 9 | Flying group members | The identities of the birds joining in each flying group, scored to account for pair and family movement together. Not always exhaustive in big groups, but focal, mate and family members were always identified. (ID). |

### Statistical Analysis

Observations with missing values for any of the variables 1 – 6 were discarded (total = 64 observations of the originally collected 651), resulting in 587 observations for inclusion in the statistical analyses. The excluded observations were the flying groups where the signalling initiator (focal) was not clearly determined (N = 52), and those where the focal was an unsexed juvenile (N = 12). All 587 observations kept for analyses included Locomotion calls, while 505 of them involved the use of the Pre-flight call. The total number of focal birds was 87 (F = 34, M = 53). For the successful flights, 13 observations consisted of a solitary flier (Flying group size = 1), and in 11 of those observations, the flier was the focal birds.

In preparation for fitting the models, we inspected the distribution of all quantitative predictors for outliers using boxplots. We removed one observation where the duration represented an extreme outlier (1200 seconds) as it was more than double the next longest duration (390 seconds), and represented an extreme case of extended signalling not observed otherwise. The continuous predictors, accounting for the number of callers in each call type, were z-scored prior to running the analyses for an easier comparison of the results.

All statistical analyses were performed in RStudio version 4.2.2 (Posit team, 2025). To investigate the factors influencing successful flight initiation and the size of the flying group, we fitted a hurdle Generalized Linear Mixed Model (GLMM; Mullahy, 1986) with a negative binomial error distribution (Lindén & Mäntyniemi, 2011), using the function *glmmTMB* from the package glmmTMB (Brooks et al., 2017). Hurdle models are two-component models that are suitable for count data containing both zero and positive values. The hurdle (zero) component models the probability that a flight initiation attempt fails, corresponding in our data to observations with a group size of zero. The conditional component models the size of the flying group conditional on successful flight initiation (group size > 0). Alternative hurdle models assuming Poisson and negative binomial count distributions were compared using Akaike’s Information Criterion (AIC) and Bayesian Information Criterion (BIC). The negative binomial model performed consistently better, and was therefore retained for all subsequent analyses.

To define our hypothesised causal pathway of the flight initiation signalling components, we considered the temporal sequence at which each component occurs (Black, 1988; Raveling, 1969; Schmitt, 1991); we decided that the sex and breeding status of the focal bird occupy the highest level on our hypothesised causal hierarchy due to their precedence in the signalling process. Second in causality are the vocal signals of interest; we decided that only the Locomotion call, expressed as the number of callers, is higher in causality because it is the first component of vocal signalling to occur (along with the posture related and motor components). At the next level of causality, we decided to include the vocal response of flock mates and family members, as their occurrence is contingent on the characteristics of the first caller and their signalling. An additional component at this level of causality is the production of the Pre-flight call by flock mates, which is not required for successful flight initiation but is observed in the majority of flight initiation attempts. The final components of the pathway represent the outcome of signalling in terms of a successful/unsuccessful flight initiation, and in the case of success, followed by flying group size. This hypothesised causal pathway structure was formalised as a Directed Acyclic Graph (DAG; Figure 3), and used to guide the selection of explanatory variables included in the statistical models. Accordingly, our model’s fixed effects included the number of responders with the Pre-flight call, the number of responders with the Locomotion call, the breeding status and sex of the focal bird, and whether all family members of the focal bird responded vocally (variables 1–5, Table 1).

**Figure 3.**
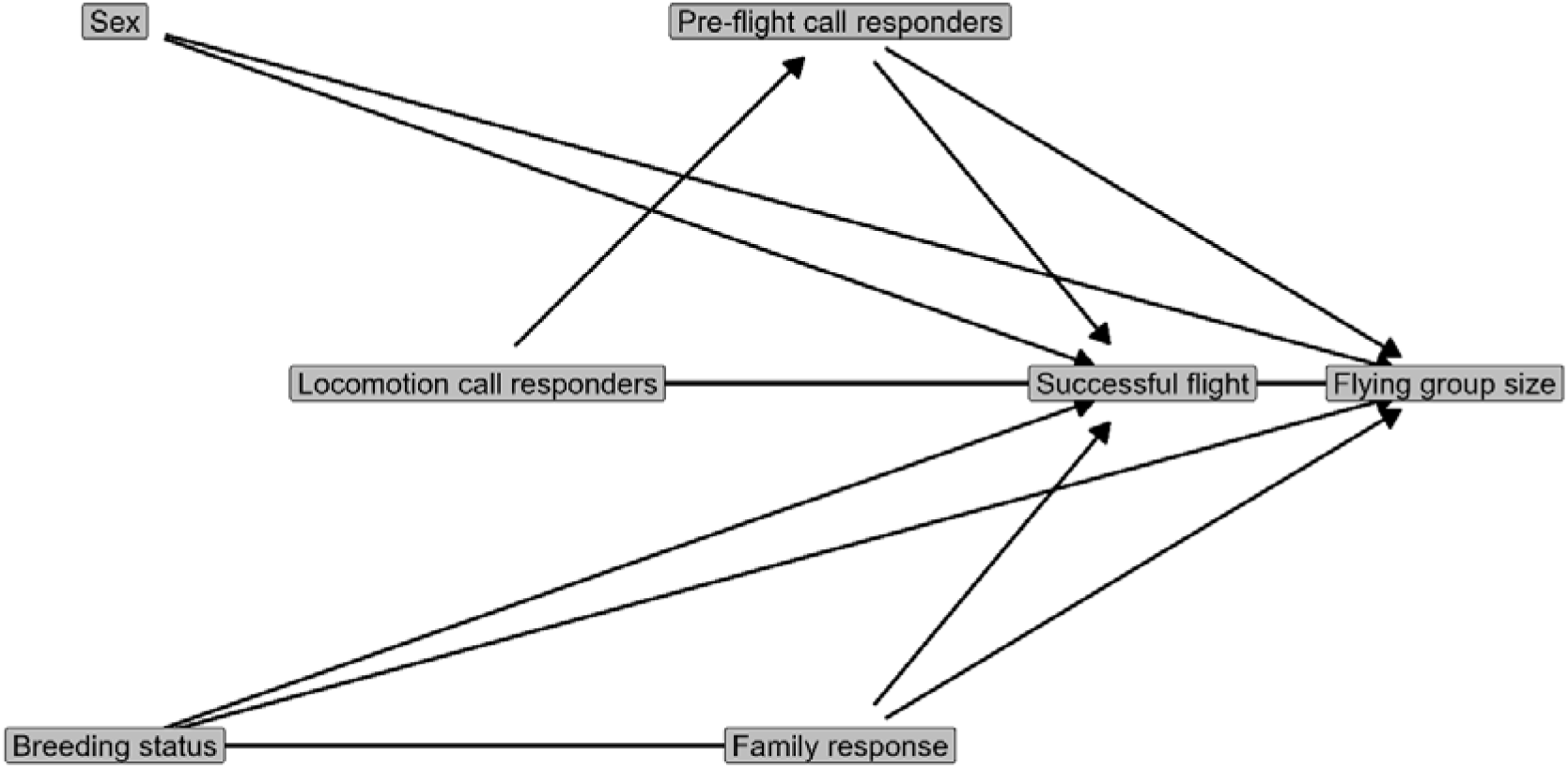
Directed acyclic graph illustrating our hypothesised causal relationships among the components of flight initiation in greylag geese. Successful flight represents whether a flight initiation attempt resulted in take-off (i.e., successful vs unsuccessful initiation), while flying group size represents the number of individuals recruited into the group. Alt text: graphical representation of the hypothesised relationship between the variables involved in the movement process under study.

To account for the non-independence of repeated observations, we initially fitted crossed random intercepts for focal identity (variable 6) and data collection session (variable 7). The random intercept for focal identity accounted for repeated observations of the same initiator, whereas the random intercept for session accounted for variation among different observation sessions, each of which comprised multiple departure attempts occurring within the same day and time period. Model diagnostics indicated that the focal random intercept in the hurdle component was estimated with a variance approaching zero, resulting in a singular fit. Consequently, the focal random intercept was removed from the hurdle component, while the session random intercept was retained. The final random-effects structure therefore consisted of (1 | Focal) + (1 | Session) for the conditional model and (1 | Session) for the hurdle component.

The significance of the conditional component was assessed using Type II Wald χ² tests implemented in the function Anova of the package car (Fox & Weisberg, 2011). Because equivalent Type II tests are not implemented for the hurdle component, the significance of each hurdle predictor was evaluated by sequentially removing individual terms from the hurdle component and comparing the models fitted with the full data set and that fitted with the subset minus outliers using likelihood ratio tests. Overall model significance was assessed by comparing the full model with a corresponding null model containing only the random effects.

Model assumptions were evaluated using simulated residual diagnostics implemented in the package DHARMa (Hartig, 2020). Residual dispersion, zero inflation and residual uniformity were assessed using the functions *testDispersion, testZeroInflation*, and *testUniformity*, respectively. Additional model diagnostics included tests for overdispersion, singularity and multicollinearity using the functions *check_overdispersion, check_singularity*, and *check_collinearity* from the package performance (Lüdecke et al., 2021). Multicollinearity among fixed effects was low, with all variance inflation factors below 1.5 (Quinn & Keough, 2002). Model stability was evaluated by sequentially removing levels of each random effect grouping factor and refitting the model to determine whether omission of individual focal identities or sessions substantially altered the parameter estimates.

We calculated 95% confidence intervals for all fixed-effect estimates using profile likelihood estimation implemented by the function *confint* in glmmTMB. Unlike Wald confidence intervals, profile likelihood intervals estimate parameter uncertainty directly from the likelihood surface and do not rely on the assumption of symmetric normally distributed sampling errors. Model coefficients and confidence intervals were reported on the link scale, while predicted values and associated 95% confidence intervals were back-transformed to the response scale for graphical presentation.

To assess whether extreme values of the responder-count variables influenced model inference (Pre-flight responders mean = 1.4, median = 1, range 0 – 15; Locomotion responders mean = 3.6, median = 3, range 0 – 20), we refit the hurdle model using a dataset in which observations above the upper Tukey cut-off (1.5 × IQR above the third quartile) for two variables were sequentially excluded. The model retained the same structure as the model fitted with the full dataset, and the estimation procedure, diagnostic checks, and inferential approach as the full model were also identical. For the final analysis, we utilised the full dataset to preserve the naturally observed distribution of the variables. The detailed modelling method and results of the model fitted with the subset dataset are presented in the Supplementary Material (Table S1; Figure S1).

Following the inspection of the model results, we ran an additional post-hoc model, aimed at investigating whether the positive effect of breeding status on flying group size was purely a consequence of the number of juveniles following their focal parent, or whether parents also recruited more non-family group members. For this analysis we selected only flight initiation attempts (N = 483), because in unsuccessful attempts, the focal bird cannot recruit non-family members, and therefore the zeros resulting from those attempts are different from those resulting from successful attempts where the number of non-family group members was zero. We fitted a negative binomial generalised linear model (GLM) with log link (McCullagh and Nelder,1989), using the function *glmmTMB* from the package glmmTMB. We used the number of only non-family members that joined in the flight group (‘Non family recruited’, variable 9, Table 1), and for predictors, we used a fixed effects structure similar to the conditional component of the hurdle model; fixed effects (variables 1, 2, 4 and 5; Table 1), but excluded the predictor ‘All family vocal response’ (variable 3; Table1) because this variable represents a downstream component of the Locomotion process and therefore is not on an equal level of causality on our hypothesised pathway (DAG 1). We omitted the inclusion of random effects following the exploration of their variance, which we found to be close to zero (<0.00001), and because their inclusion resulted in model singularity. Comparison of models with and without random intercepts produced identical fixed-effect estimates, but the simpler model showed lower AIC and BIC values. Consequently, the final model was fitted without random effects.

Model diagnostics, and assumptions were checked using the same tests utilised for the hurdle model, and model significance, confidence intervals, and back transformation of the estimates to the response scale for reporting were also performed using the same functions utilised for the hurdle model. For the model interpretations in the results section, the model estimates were back transformed from the model link scale to the response scale by exponentiation. Estimates of the hurdle/model component (logit link) were exponentiated to obtain odds ratio of unsuccessful flight initiation, and the estimates of the conditional component (log-link) were exponentiated to obtain rate ratios for expected flying group sizes. All estimate interpretations are reported in the results section as percentage changes in the odds of unsuccessful flight initiation and expected flying group sizes, respectively, for each one standard deviation increase in the continuous predictors, or relative to the reference level of the categorical predictors. Model predictions presented in the figures were back transformed to the response scale and are shown as the probability of successful flight initiation for the hurdle/zero component, and predicted flying group sizes for the conditional component.

## Results

Among the 587 observations of flight initiation attempts, 483 (82%) resulted in successful flight initiation. Out of the total number of trials, 195 (33%) observations had a focal bird in a pair with juveniles, and 342 (58%) had a focal bird in a pair without any juvenile. The remaining 22 (3.7%) observations had a focal bird that was not paired (sub-adults, and senior males). In 93% of the trial where the focal bird had juveniles, the family stayed together (flew together in case of success and none departed for aborted attempts). For pair-bonded focal birds without juveniles, the percentage of pairs remaining together was 97%.

### Probability of successful flight initiation

The full hurdle model fitted the data significantly better than the null model, indicating a significant contribution of the fixed effects towards explaining the variance of the data (Likelihood ratio test: χ²₁₀ = 171.07, P < 0.0001). The hurdle (zero) component, investigating the probability of unsuccessful flight initiation (Group size = 0) revealed that, at the mean values of the continuous predictors and at the reference levels of the categorical ones, the estimated probability of successful flight initiation was 81.4 %.

The vocal response of all family members had the strongest positive effect on the probability of successful flight initiation, increasing the odds of successful flight initiation by 68% relative to the response of some or none of the family members (Figure 4 C, Table 2). One standard deviation increase in the number of Locomotion responders reduced the odds of failure by approximately 45% (Figure 4 A), while every standard deviation increase in the number of Pre-flight responders reduced those odds by approximately 34% (Figure 4 B, Table 2).

**Figure 4.**
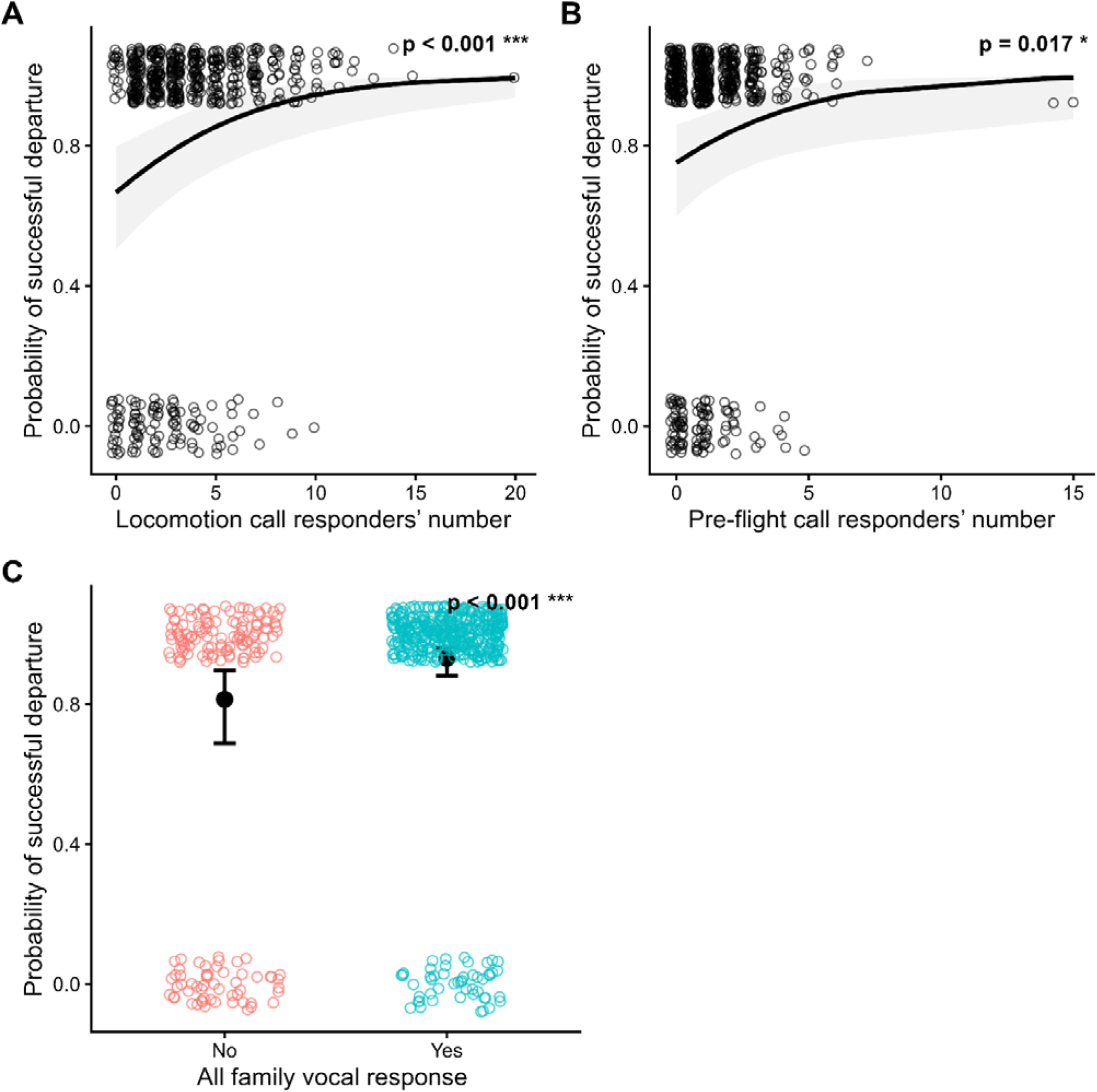
Predictors influencing the successful flight initiation of greylag geese. Hurdle model – zero component: effects of the model predictors on the probability of successful flight initiation; (A) Effect of the number of birds participating in Pre-flight calling during the trial. (B) Effect of the number of birds participating in Locomotion calling during the trial. (C) Effect of the vocal participation of all family members in Pre-flight calling (No vs. Yes). Predictions are presented on the response scale. Data points represent the distribution of the observed values, the trend line shows the predicted fit, and the shaded area for A and B plus the error bars for C represent model 95% confidence intervals. The continuous predictors were standardised (z-scored) prior to modelling. Alt text: graphical representation of the first model significant findings. Three panels title a, b, and c are shown in order, each representing the relationship between the first response variable and three significant predictors

**Table 2.** Model results investigating the variables influencing successful flight initiation and flying group size in greylag geese group flight. Parameter estimates for the conditional (group size) and hurdle/zero (probability of successful flight initiation) components of the hurdle negative-binomial GLMM. Continuous predictor variables (Pre-flight callers and Locomotion callers) were z-scored before analysis. Regression coefficients and profile-likelihood 95% confidence intervals are presented on the model link scale (logit link for the hurdle/zero component and log link for the conditional component). The conditional component significance was assessed using Type II Wald χ² tests, while the hurdle component significance was assessed by likelihood-ratio tests comparing models with and without each predictor. The negative coefficients in the zero component indicate a decrease in the probability of zero (unsuccessful flight initiation).

| **Zero component (Successful flight initiation)** |  |  |  |  |
| --- | --- | --- | --- | --- |
| **Predictor** | **Estimate ± SE** | **95% CI** | **$\chi^2$** | **P** |
| (Intercept) | - 1.474 ± 0.349 | -2.188 – -0.814 | | **< 0.0001\*\*\*** |
| Locomotion call responders' number | -0.597 ± 0.172 | -0.949 – -0.275 | 14.160 | **< 0.0001\*\*\*** |
| Pre-flight call responders' number | - 0.411 ± 0.173 | -0.766 – -0.086 | 6.292 | **0.017 \*** |
| Breeding status (parent) | 0.386 ± 0.283 | -0.169 – 0.942 | 1.865 | 0.172 |
| Sex (Male) | 0.314 ± 0.280 | -0.225 – 0.878 | 1.290 | 0.261 |
| All family vocal response (Yes) | - 1.130 ± 0.261 | -1.646 – -0.619 | 18.590 | **< 0.0001\*\*\*** |
| **Conditional component (Group size)** |  |  |  |  |
| **Predictor** | **Estimate ± SE** | **95% CI** | **$\chi^2$** | **P** |
| (Intercept) | 1.727 ± 0.121 | 1.485 – 1.964 | | **< 0.0001\*\*\*** |
| Locomotion call responders' number | 0.329 ± 0.039 | 0.253 – 0.408 | 69.620 | **< 0.0001\*\*\*** |
| Pre-flight call responders' number | 0.073 ± 0.035 | 0.004 – 0.144 | 4.185 | **0.041 \*** |
| Breeding status (parent) | 0.308 ± 0.114 | 0.091 – 0.550 | 7.250 | **0.007 \*\*** |
| Sex (Male) | 0.032 ± 0.087 | -0.173 – 0.240 | 0.096 | 0.756 |
| All family vocal response (Yes) | - 0.143 ± 0.087 | -0.316 – 0.028 | 2.684 | 0.101 |

The conditional component (conditional on the success of flight initiation, Group size > 0), revealed that, at the mean value of all continuous predictors, and when categorical predictors were at their reference level, the estimated flying group size was 5.6 individuals. Flying group size increased most significantly with the increase of Locomotion call responders, with each standard deviation increase corresponding to 39 % increase in expected flying group size (Figure 5 A; Table 2). The increase in Pre-flight call responders also had a positive effect on flying group size, with each standard deviation increase corresponding to 7.5 % increase in expected flying group size (Figure 5 B), while parents departed with expected flying groups approximately 36 % larger than non-parents (Figure 5 C; Table 2).

**Figure 5.**
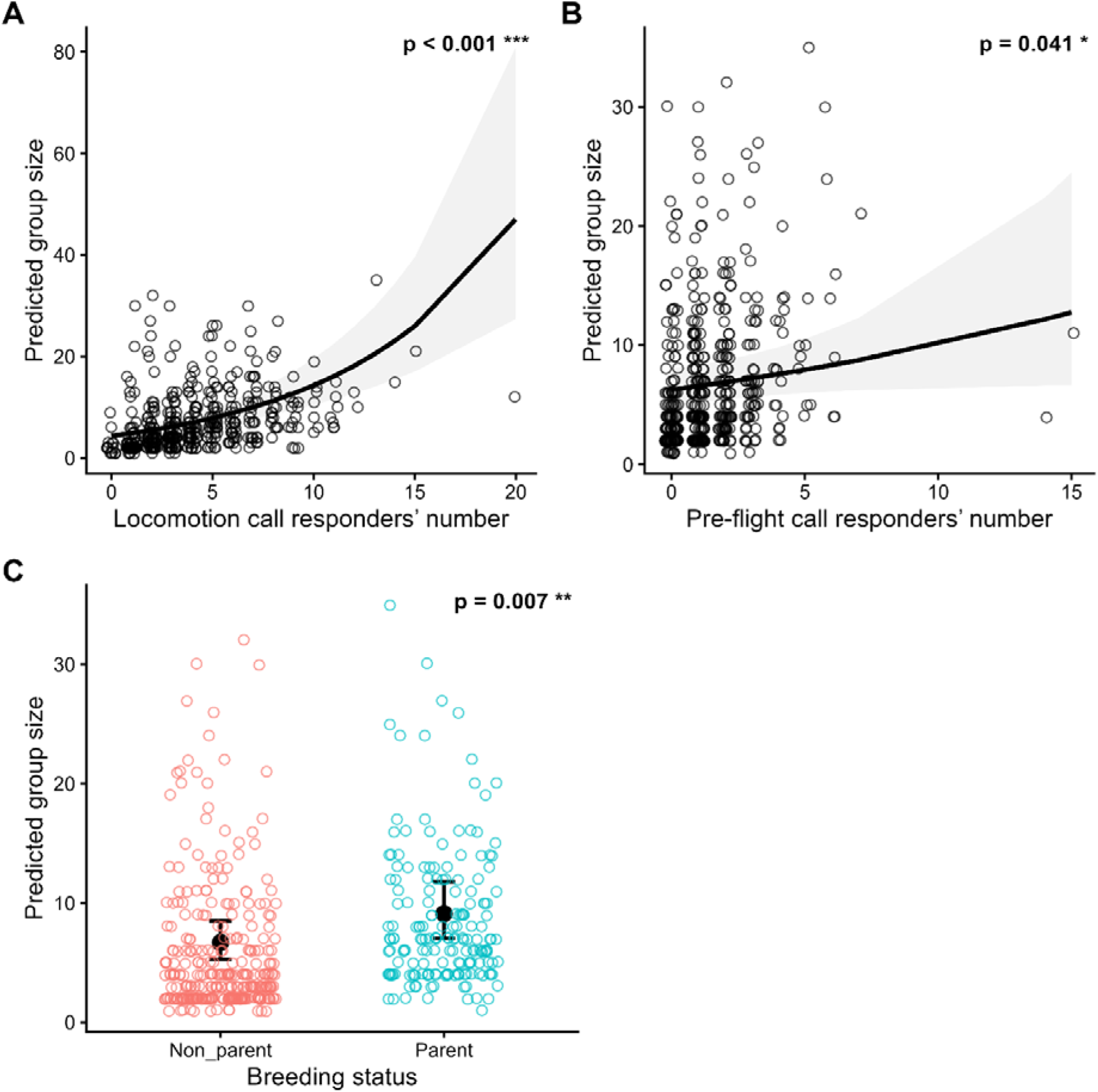
Predictors influencing the flying groups size of greylag geese collective flights. Hurdle model – conditional component: effects of the model predictors on the flying group size; (A) Effect of the number of birds participating in Pre-flight calling during the trial. (B) Effect of the number of birds participating in Locomotion calling during the trial. (C) Effect of the breeding status of the focal bird (Parent / non-parent). Predictions are presented on the response scale. Data points represent the distribution of the observed values, the trend line shows the predicted fit, and the shaded area for A, and B plus the error bars for C represent model 95% confidence intervals. The continuous predictors were standardised (z-scored) prior to modelling. Alt text: graphical representation of the first model significant findings. Three panels title a, b, and c are shown in order, each representing the relationship between the second response variable and three significant predictors

The model fitted with subset data showed the same terms to be significant (Table S1, Supp. Mat.), and the estimates for the model did not deviate considerably from those of the full model (Table 2, Table S1). Overall, an increase in the number of vocal responders – using both call types – predicted an increase in the probability of successful flight as well as an increase in flying group size. The vocal response of all family members of the focal bird predicted an increase in successful flight initiation, while parents departed with larger groups compared to non-parents.

### Do parent groups attract more non-related birds for flight

The full model investigating the effect of breeding status, number of responders in both call types, and the sex of the focal bird on the number of non-family members joining in departure group provided a better fit for the data compared to the null model; the full-null model comparison revealed a significant contribution of the fixed effects towards explaining the variance of the data (Likelihood ratio test: χ²_4_ = 39.93, P < 0.0001).

The only significant predictor of the increased number of non-family members recruited in flying groups was the number of Locomotion call responders (Table 3, Figure 6).

**Figure 6.**
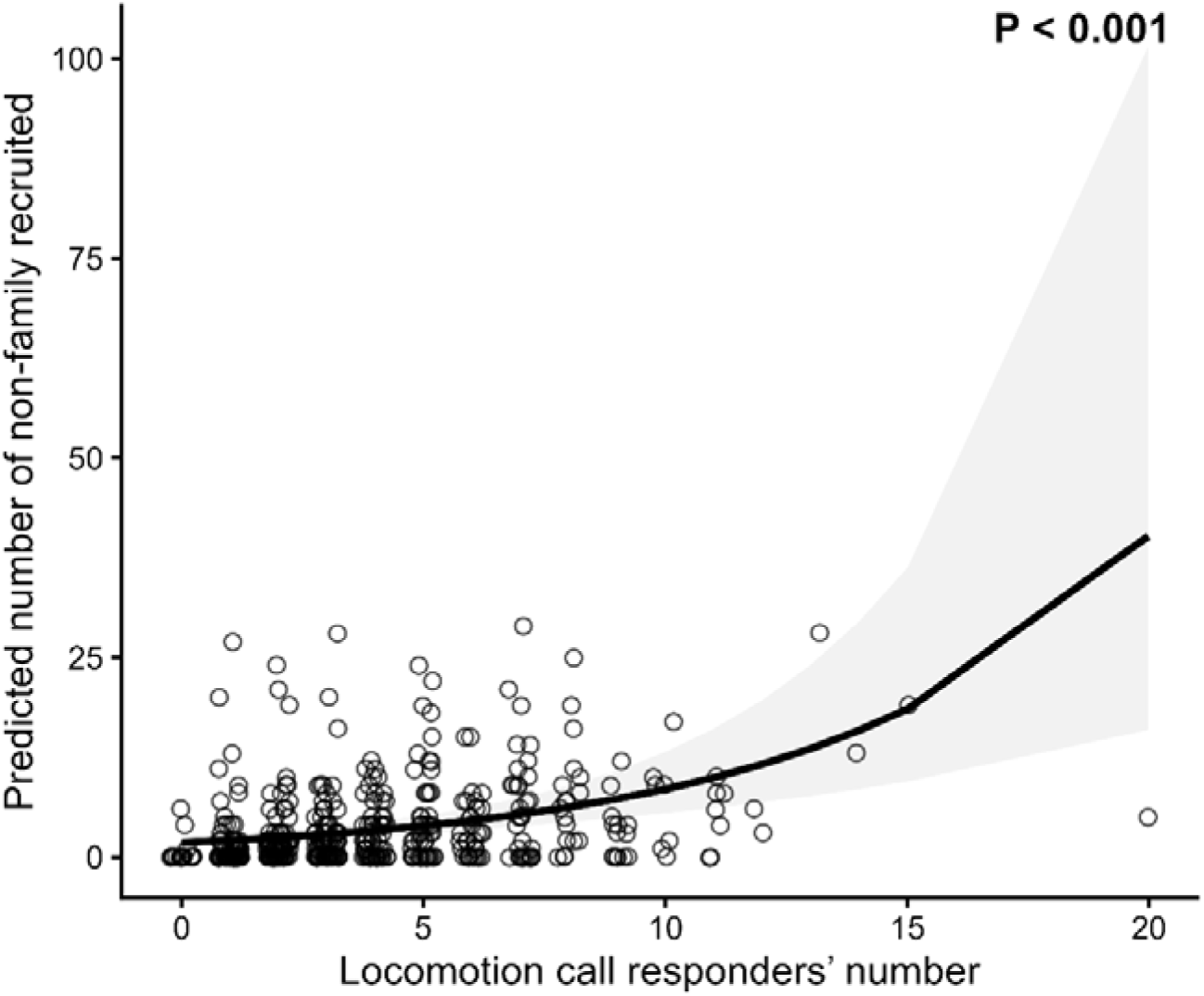
Predictors that significantly influenced the number of non-family members recruited to greylag geese collective flights. Effect of the number of Locomotion call responders on the predicted number of non-family individuals recruited into the departure group. Data points represent the distribution of the observed values, the trend line shows the predicted fit, and the shaded area represents model 95% confidence intervals. Predictions are presented on the response scale. The number of Locomotion call responders was standardised (z-scored) prior to modelling. Alt text: graphical representation of the second model significant finding, representing the relationship between the response variable and the only significant predictor.

**Table 3.**
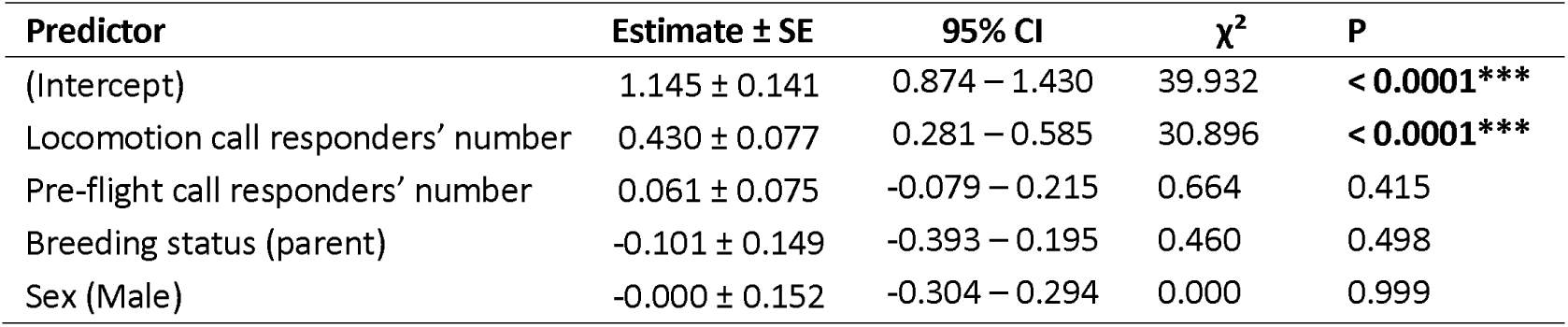
Predictors influencing the recruitment of non-family members to flying groups in greylag geese collective flight. Parameter estimates for the negative-binomial GLM investigating the effect of the predictors on the number of non-family individuals joining in flying group. Continuous predictor variables (Pre-flight responders and Locomotion responders) were z-scored before analysis. Regression coefficients and profile-likelihood 95% confidence intervals are presented on the model link scale (log link). Parameter significance was assessed using Type II Wald χ² tests.

| Predictor | Estimate ± SE | 95% CI | $\chi^2$ | P |
| --- | --- | --- | --- | --- |
| (Intercept) | 1.145 ± 0.141 | 0.874 – 1.430 | 39.932 | <0.0001*** |
| Locomotion call responders' number | 0.430 ± 0.077 | 0.281 – 0.585 | 30.896 | <0.0001*** |
| Pre-flight call responders' number | 0.061 ± 0.075 | -0.079 – 0.215 | 0.664 | 0.415 |
| Breeding status (parent) | -0.101 ± 0.149 | -0.393 – 0.195 | 0.460 | 0.498 |
| Sex (Male) | -0.000 ± 0.152 | -0.304 – 0.294 | 0.000 | 0.999 |

Each increase of one standard deviation in the number of Locomotion call responders corresponded to a 54 % increase in the number of non-family individuals recruited in the flying group (rate ratio = 1.54, 95% CI = 1.33–1.80). In contrast to the full model, the positive relationship between parent breeding status and the increase of flying group size seen in the conditional component of the hurdle model disappeared when the response was restricted to non-family members joining in the flying group of the focal bird (Table 3).

## Discussion

Greylag geese groups move in cohesive flocks during autumn and winter, and the flock might break into smaller flying groups of various sizes that coalesce on land (Lorenz, 1988; Schmitt, 1991). The species utilises specialised calls to initiate spontaneous group flight (Cramp & Simmons, 1977). Because of their long-lasting social bonds, it is expected that the association of greylag geese during flight is non-random, and that the decision to join others in flight is therefore influenced by information contained in their flight initiation calls (Mostafa & Briefer, 2026). Through observing spontaneous flock movement between different sites during autumn, we investigated how these birds utilise flight initiation calls to initiate group flight, and recruit others into their flight group. Our findings suggest that both the identities of the birds participating in flight initiation calling, as well as their numbers, influence the decision of the signalling goose to either fly or abort the flight imitation attempt. The compositions and sizes of the successfully initiated flying groups are consistent with a system where flight decisions are made both to maintain the cohesion of basic family unit, and to maximise the flight group size. The collective decision to fly in greylag geese is thus a combined decision at the level of the individual (Conradt & Roper, 2005), where knowledge about own motivation and readiness to fly are integrated with public information shared by the group, including both family members and flock mates, to make the decision of whether to join a flying group.

As predicted, vocal signals played an essential role in the flight initiation signalling of the species; all flight initiation attempts that we observed were preceded by the stereotyped signalling described in geese (Raveling, 1969; Schmitt, 1991), which consistently incorporates at least the Locomotion call. This call was emitted in our study by all the identified flying individuals, confirming the vocal signalling pattern reported in an earlier study (Schmitt, 1991). The use of context-specific vocal signals to coordinate the timing of the onset of movement is widespread among species of high group cohesion. For example, the green woodhoopoes *Phoeniculus purpureus*, a social species that inhabits thick forest habitat and maintain cohesive groups, relies entirely on vocal coordination for moving between foraging sites (Radford, 2004). Similarly, movement coordinate in groups spread over large forging areas is achieved via the use of vocal signals, bypassing the need to keep group mates in sight (Stewart & Harcourt, 1994). The use of flight initiation calls that are produced in a chorus is similarly conserved in related species including Canada geese, where flocking individuals join in calling and head tossing prior to flying in smaller, consecutive groups (Raveling, 1969). Pre-flight calls, on the other hand, occurred in the majority – but not all – of our observations closer to take-off. This call type was emitted by a subset of the signalling individuals if at all, suggesting that the two calls call types have different functions within the flight coordination process, as typically found in calls with different audible ranges produced in the same context (Naguib & Wiley, 2001; Titze & Palaparthi, 2018; Liao et al., 2024).

The probability of flight success increased with the number of individuals participating in flight initiation calling, irrespective of the call used, but the number of birds producing Locomotion calls had a larger and more robust effect on flying group size. Similarly, larger flying group sizes were observed when more callers joined in flight initiation calling using either call type, with the number of Locomotion callers having a larger and more robust effect on the resulting flying group size. This effect is also observed in the collective movement initiation of other species with similar ecological needs of foraging and protection, including domestic geese *Anser domesticus* (Ramseyer et al., 2009), where a higher number of vocalising individuals and increased arousal prior to movement predicts larger groups during movement. The consistent and considerable effect the number of Locomotion callers had on both the successful initiation of flight and recruiting others in our study is consistent with systems where a threshold of callers needs to be reached before a collective decision is reached, whether a minimum number or percentage of participants achieves quorum (C. Wang et al., 2020), or where the number of callers required to reach quorum depends on the identity of the initiator (Walker et al., 2017). It is worth mentioning that the ability to initiate successful flight did not seem to vary among individuals, while flying group size did, further suggesting that the individuals differ in their ability to recruit others in our study system.

We had predicted that the signalling individuals (focal birds) would make the decision to join in a flight group as a pair or a family, and that family members would always move together. Accordingly, the vocal response of the mate in the case of pair-bonded individuals, and that of the mate and offspring in the case of families, was the predictor with the largest positive influence on the probability of a successful flight. This relationship illustrates that individuals maintain proximity to mates and family on the move. Similarly, the coordination of group movement in multiple monkey species is influenced by the affiliative relationships and kinship of the individuals joining in moving groups (Jacobs et al., 2011; Seltmann et al., 2013; Sueur, Deneubourg, et al., 2011). Maintaining proximity to social affiliates, mate and offspring is necessary for mate guarding, parental care, and shared support during conflict (Frigerio et al., 2001). Pair-bonded individuals of some large waterfowl species show increased fitness by maintaining proximity to offspring (Black & Owen, 1989), as well as proximity to the mate, since lifetime reproductive success decreases with re-pairing (Black, 2001). Juvenile greylags are dependent on their parents for the first year of their lives (Lorenz, 1988; Rutschke, 1982), and during that year, the parents provide constant parental care in the form of protection, warding off conspecifics during foraging and fighting off aggressors (Scheiber et al., 2005; Szipl et al., 2019). In migratory populations, the first autumn of a juvenile’s life also coincides with migration (Andersson et al., 2001), during which the juveniles travel through unfamiliar terrain, making family cohesion and proximity to experienced parents particularly vital for survival of the offspring and subsequently for the fitness of the parent. The effect of mate and family member vocal response on the probability of a successful flight that we highlighted is consistent with our findings that the flight initiation calls of greylag geese carry individual signature (Mostafa & Briefer, 2026), and that paired individuals respond to the broadcast of partner locomotion calls at higher rates compared to calls of other familiar conspecifics (Mostafa & Briefer, in preparation). This is likely highly useful as signalling geese do not always keep mate and offspring in line of sight until they fly because everyone faces the same direction of intended flight (Schmitt, 1991). The positive influence of mate and offspring on the probability fo successful flight is also consistent with a decision-making mechanism where signals are identity-weighted, and those of social affiliates contribute a larger effect towards reaching the decision to move (Marshall et al., 2017)

We found a positive relationship between the presence of juveniles and the flying group sizes that included family members (Conditional component, Table 2), but it had no effect on the recruitment of non-family members (Table 3). The effect seen in the conditional component of the hurdle model, where parents recruited larger group sizes, can be interpreted in the light of families staying together 94% of the time in our sample; when a parent recruited only their family, the smallest flying group consisted of three individuals, while a non-parent recruiting only their mate flew in a group of two. As a result, parents with juveniles always recruited larger flying groups on average. The results of the additional model investigating the factors that influenced the recruitment of non-family members (Table 3), found that only the number of Locomotion callers was significantly associated to bigger flying group sizes. This finding provides insight into the result of the conditional component of the hurdle model; the vocal participation of juveniles in flight initiation signalling started by a parent result in a higher number of callers compared to the attempts of non-parents, where only the vocal response of mate is almost guaranteed. The increased number of Locomotion callers in parent attempts should therefore result in increased probability of reaching the quorum necessary for departure, and in a shorter time. Therefore, it is reasonable to conclude that the presence of juveniles helps to amplify their family’s signal, a common phenomenon of positive feedback, where mimetism - defined as the process by which the higher number of individuals performing a behaviour, the higher the probability that others will join in the behaviour – is a contributing mechanism to the decision making process (Sueur et al., 2011; Sueur & Deneubourg, 2011).

Combining these results starts to elucidate the first elements involved in the group flight decisions of greylag geese; the significance of the number of callers along with their identities – family vs. non family – suggest a quorum mechanism with identity-weighted signals, while the success of parents in recruiting larger groups, including their mate and offspring, is consistent with a system where mimetism plays some role in reaching the threshold required for flight. Further downstream variables not investigated in this study, including the temporal elements, could also affect the final decision-making mechanism in the species.

## Conclusion

Our findings are consistent with a system where the decision to fly and to join others in flight is a combined decision. Our results illustrate how the flight initiation signalling in greylag geese can inform individual of the number and identities of callers, allowing them to join the flight group which maintains the cohesion of the mate and family units, while also maximizing the benefit of staying in larger groups. We expect closely-related species with similar patterns of social organisation to utilise similar strategies of flight coordination, employing specialised signals for local and group-wide coordination, and to also prioritise family cohesion and larger flying group sizes. Further investigation of the specific function of each call type could shed a light on the reason behind the difference in their timing and their audible range, as well as the possible syntactic meaning of their production in tandem. Experimental investigation of possible interaction between the identities of the callers and the resulting departure group size is of interest to researchers working on leader-follower dynamics.

## Supporting information

Supplemental modelling results

## Acknowledgements

We thank Konrad Lorenz Research Centre for Behaviour and Cognition in Grünau im Almtal (47_48050.500N 13_56051.000E), and Professor Sonia Kleindorfer for making this work possible by providing accommodation and access to the banded Greylag goose population during three field seasons, as well as making all the equipment necessary for data collection available. We would also like to extend thanks to Dr. Didone Frigerio and Dr. Josef Hemetsberger for access to the long-term life history data of the goose population and for generously making their invaluable knowledge of the population and the species available. We would like to thank Ariana Strandburgh-Peskin for her expert input and comments on this manuscript, following many discussions of the theory and experimental design, Professor David Richard Nash for his feedback and discussion of the statistical analyses and their interpretation, and Professor Tecumseh Fitch for providing extensive commentary and valuable edits to the manuscript.

This study was partially supported by a Semper Ardens: Accelerate Fellowship from the Carlsberg Foundation awarded to E.F.B. (CF20-0538).

## Data Availability

https://doi.org/10.5281/zenodo.17691992

## Author contribution

D.A.A.M.: Conceptualisation, Project administration, Investigation, Data curation, Formal analysis, Methodology, Writing – original draft.

E.M.B.: Supervision, Validation, Writing – review & editing.

N.M.A.: Validation, Writing – review & editing

A. G. M.: Writing – review & editing.

## Notes

https://doi.org/10.5281/zenodo.17691992

### Competing Interest Statement

The authors have declared no competing interest.

