## Supplemental modelling results for "Greylag geese (Anser anser) initiate group flight based on the number and identities of callers"

**Supplementary Material**

***Model fitted with data minus outliers to investigate whether the outliers of the continuous predictors affect model inference:***

The results of the hurdle model (Table S1; Figure S1), fitted with the data minus the upper outliers, revealed parameter estimates consistent with those of the model fitted using the full dataset (Table 2). The only notable difference was that the positive association between the number of departure responders and departure group size in the conditional component was no longer statistically significant, while all other significant estimates remained unchanged.

Table S1– Model results investigating the variables influencing successful flight initiation and flying group size in greylag geese group flight, fitted with data minus the upper outliers of the continuous predictors. Parameter estimates for the hurdle model, consisting of the hurdle/zero (probability of successful flight initiation) and the conditional (group size) components of the hurdle negative-binomial GLMM. Continuous predictor variables (departure responders and recruitment responders) were z-scored before analysis. Regression coefficients and profile-likelihood 95% confidence intervals are presented on the model link scale (log link for the conditional component and logit link for the hurdle component). The conditional component significance was assessed using Type II Wald χ² tests, while the hurdle component significance was assessed by likelihood-ratio tests comparing models with and without each predictor. The negative coefficients in the zero component indicate a decrease in the probability of zero (unsuccessful flight initiation)

| **Conditional component** | |  | |  | |  | |
| --- | --- | --- | --- | --- | --- | --- | --- |
| **Predictor** | **Estimate ± SE** | | **95% CI** | | **χ²** | | **P** |
| (Intercept) | 1.733 ± 0.128 | | 1.478 – 1.983 | |  | | **< 0.0001***** |
| Pre-flight call responders' number | 0.070 ± 0.051 | | -0.030 – 0.171 | | 1.870 | | 0.172 |
| Locomotion call responders' number | 0.394 ± 0.051 | | 0.295 – 0.496 | | 59.786 | | **< 0.0001***** |
| Breeding status (parent) | 0.314 ± 0.122 | | 0.083 – 0.571 | | 6.610 | | **0.010*** |
| Sex (Male) | -0.001 ± 0.107 | | -0.214 – 0.213 | | 0.000 | | 0.990 |
| All family vocal response (Yes) | -0.129 ± 0.093 | | -0.311 – 0.053 | | 1.937 | | 0.164 |
| **Zero component** | | |  | |  | |  |
| **Predictor** | **Estimate ± SE** | | **95% CI** | | **χ²** | | **P** |
| (Intercept) | -1.485 ± 0.357 | | -2.215 – -0.808 | |  | | **< 0.0001***** |
| Pre-flight call responders' number | -0.383 ± 0.186 | | -0.761 – -0.029 | | 4.518 | | **0.034*** |
| Locomotion call responders' number | -0.637 ± 0.187 | | -1.017 – -0.283 | | 13.071 | | **< 0.0001***** |
| Breeding status (parent) | 0.432 ± 0.287 | | -0.131 – 0.997 | | 2.266 | | 0.132 |
| Sex (Male) | 0.306 ± 0.282 | | -0.236 – 0.873 | | 1.205 | | 0.272 |
| All family vocal response (Yes) | -1.140 ± 0.263 | | -1.659 – -0.625 | | 18.581 | | **< 0.0001***** |

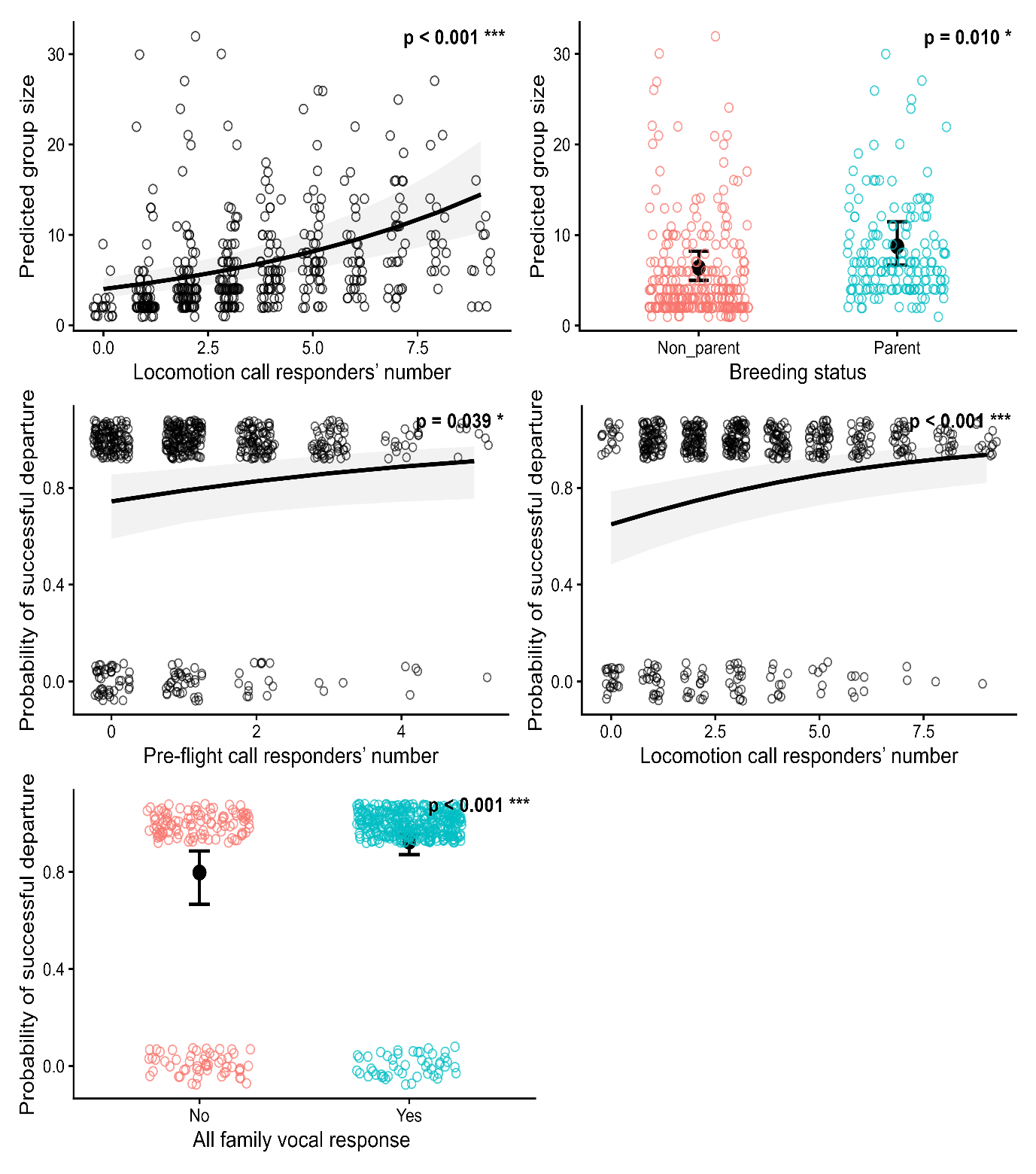

Figure S1 – Predictors influencing the successful flight initiation of greylag geese. Hurdle model fitted with data minus continuous predictors upper outliers – zero component: effects of the model predictors on the probability of successful flight initiation; (A) Effect of the number of birds participating in departure calling during the trial. (B) Effect of the number of birds participating in recruitment calling during the trial. (C) Effect of the vocal participation of all family members in pre-flight calling (No vs. Yes). Predictions are presented on the response scale. Data points represent the distribution of the observed values, the trend line shows the predicted fit, and the shaded area for A, and B plus the error bars for C represent model 95% confidence intervals. The continuous predictors were standardised (z-scored) prior to modelling.

Alt text: graphical representation of the significant findings for the hurdle model fitted with the data minus the outliers of the two continuous predictors. Five panels are shown in order, the first two representing the relationship between one response variable and two significant predictors, while the latter three represent the relationship between the other response variable and three significant predictors
